# Analysis of the Quality of Macromolecular Structures

**DOI:** 10.1101/108456

**Authors:** Dinakar M. Salunke

**Affiliations:** International Centre for Genetic Engineering and Biotechnology Aruna Asaf Ali Marg, New Delhi-110 067. India.

## Abstract

Structure determination utilizing X-ray crystallography involves collection of diffraction data, determination of initial phases followed by iterative rounds of model building and crystallographic refinement to improve the phases and minimize the differences between calculated and observed structure factors. At each of these stages, a variety of statistical filters exist to ensure appropriate validation. Biologically important observations often come from interpretations of signals that need to be carefully deciphered from noise and therefore human intervention is as important as the automated filters. Currently, all structural data are deposited in the Protein Data Bank and this repository is continuously evolving to incorporate possible new improvements in macromolecular crystallography. The journals that publish data arising from structural studies modulate their policies to take cognizance of new improved methodologies. The PDB and journals have evolved an accepted protocol to ensure the integrity of crystallographic results. As a result, the quality of available data and interpretations are becoming better over the years. However, there have been periodic efforts by some individuals who misuse validation mechanisms to selectively target published research through spurious challenges. These actions do more harm to the field of structural biology and runs counter to their claim to cleanse the system. The scientific systems in structural biology are robust and capable of self-correction and unwarranted vigilantism is counterproductive.

## Introduction

As in other fields, scientific data in biology is peer-reviewed and documented. It is available to other scientists so that new experiments could be further designed to provide next level of insights into biological processes under scrutiny. Therefore, quality control of all scientific data in biology is crucial. The structural biology community in the world has been at the forefront of the efforts to render open access to data and resources. Macromolecular crystallography involves crystallization, X-ray diffraction data collection, and determination of phases to generate an electron density map, which is used to build an initial model. This preliminary model is then subjected to iterative rounds of model building and refinement to generate a final model that can be analyzed to obtain deep insights regarding the biological function of the macromolecule(s) under scrutiny. Although the steps are straightforward, and currently blessed with a large degree of automation, a significant number of crystals are of a quality that renders structure determination, model building or crystallographic refinement a tricky exercise. To extract useful information from such datasets requires careful refinement, regular examination of the maps and a deviation from the practice of “black box” crystallography. The contents of the asymmetric unit have to be modeled appropriately using basic principles of protein structure and physical chemistry. This is particularly important if the molecules adopt more than one conformation. The proper estimation of the asymmetric unit contents leads to maps that are clear enough to build a model that can be used to elucidate the structural basis for molecular function.

A traditional approach to model building and crystallographic refinement is beneficial for these challenging datasets. Model building and crystallographic refinement have to be done gradually to ensure that the phases improve steadily and provide the highest probability for new features to appear in the electron density map. Even small changes in the model have to be followed by crystallographic refinement and computation of a new map. The processes of model building and refinement have to be repeated a large number of times to reduce the noise and improve the clarity in the maps. Errors in describing the contents of the asymmetric unit or excessively hasty refinement- as is done due to automation- will not give rise to a decipherable map and may lead to the erroneous judgment that the quality of data from these crystals is inadequate to enable structure determination and refinement. The process of obtaining useful information from these challenging datasets is labor intensive and different from the current practice of “black-box” crystallography where the electron density maps are examined only few times and modeling/ligand-fitting and crystallographic refinement are carried out mostly by highly automated softwares. Also, there are further considerations in the refinement of protein structures such as, those involving conformational changes in the protein that occur due to ligand binding. The observation of such structural changes is also an important factor for establishing the presence of a ligand in the structure.

Generally, an experiment in biological crystallography is carried out in the context of a hypothesis, and the data should help prove or disprove the hypothesis. It is important to highlight that most practicing macromolecular crystallographers address a problem in biology, design biochemical experiments, execute them and then carry out the crystallographic analysis. The purification of adequate amounts of the target protein to high homogeneity and subsequent crystallization requires considerable amount of effort, and the researcher has to struggle quite a lot even before reaching the crystallographic experimental stage. With regards to macromolecular complexes associated with biology and medicine, the degree of difficulty to obtain structures is further enhanced. Indeed, as more and more complex and physiologically relevant molecular systems are selected for scrutiny, the challenges faced by crystallographers’ will only amplify. The crystallographic skills are of utmost importance in dealing with these challenges.

## Parameters for Quality Analysis

### Electron Density Map

After one has determined the correct set of phases, the interpretation of the electron density map is the most critical step in terms of elucidation of macromolecular structure from diffraction data. Obviously the guiding principle is the molecular stereochemistry and if the protein sequence is known and the diffraction data are available at very high resolution, the process currently is near automatic. However, at relatively lower resolutions, the experience of a crystallographer becomes paramount. One relies hugely on the immediate chemical neighborhood of the polypeptide chain while building the model into electron density. The presence and fit of ligands in a crystal structure is judged on the basis of many inter-related factors. Several kinds of maps are routinely used to model the structure including 2*F*o-*F*c and simulated annealing 2*F*o-*F*c composite omit maps. Delete-refine *2Fo-Fc* and delete-refine *Fo-Fc* maps are used periodically for examination of the positive electron density corresponding to the ligand model being built (Bhat, 1988; Brunger et al., 1997; Read, 1986). Model building is followed by refinement and the change in *R*_free_ and R_cryst_ is continuously monitored to ensure correct modeling of the ligand and prevent any wrong fit or over-interpretation. The overall fit of the model to the electron density is evaluated by the R-factors and the local fit is assessed through careful examination of the electron density maps. The goodness of fit locally can be evaluated using the Real Space Correlation Coefficient in majority of cases. However, these calculations may be considered cautiously in special cases, such as when the bound ligand shows multiple conformations or low occupancy. The excessive dependence on any single factor or automated programs can be detrimental to an unbiased evaluation of the model fit.

Electron density maps are generally analyzed at a variety of different σ cut-offs and there aren’t any fixed rules for setting contour levels when building models in delete-refine *F*o-*F*c electron density maps or the other electron density maps. Experienced crystallographers critically evaluate results during the refinement process to ensure that the signal-to-noise distinction is adequately handled, and go back and forth at different contour levels rather than sticking to a specified σ cut-off while analyzing different electron density maps simultaneously. It is important to emphasize that many structures, which are crystallographically sound and physiologically interesting, have ligands visualized at the contour levels less than 3.0σ. In fact, the crystallography community has recognized that precisely defining the contour level for ligand building is not possible (Adams et al., 2016; Read and Kleywegt, 2009). We provide many examples from among the biologically relevant protein-ligand complexes reported with visualization of electron density at σ cut-offs of lower than 3.0 (Table 1). Figure 1 shows delete-refine *F*o-*F*c maps of ligands contoured at 3.0σ from two structures deposited in the Protein Data Bank (PDB). It is evident that neither of these structures would have seen the light of the day if they were subjected to 3.0σ cut-off criterion. Overall, the signal can be distinguished from noise at σ cut-off lower than 3.0 in many structures.

**Table 1:** Contour levels of *F*o-*F*c electron density maps for visualizing ligands in representative structures in PDB.

| Sl. No | PDB code | Resolution (Å) | $\sigma$ cut-off | Reference |
| --- | --- | --- | --- | --- |
| 1 | 3R6R | 2.4 | 1.5 | (Bazzicalupi et al., 2013) |
| 2 | 3KO4 | 2.7 | 2.5 | (Bhatt et al., 2010) |
| 3 | 3E2Y | 2.3 | 1.5 | (Han et al., 2009) |
| 4 | 2E2Z | 2.8 | 1.5 | (Han et al., 2009) |
| 5 | 2G9Z | 2.0 | 1.5 | (Santini et al., 2008) |
| 6 | 2IAJ | 2.5 | 2.0 | (Das et al., 2007) |
| 7 | 1XZ0 | 2.8 | 1.8 | (Zajonc et al., 2005) |
| 8 | 1UVI | 2.2 | 1.5 | (Salgado et al., 2004) |
| 9 | 1UVM | 2.0 | 1.5 | (Salgado et al., 2004) |
| 10 | 1R15 | 2.4 | 1.7 | (Love et al., 2004) |
| 11 | 1PWW | 2.8 | 2.0 | (Turk et al., 2004) |
| 12 | 1UJJ | 2.6 | 1.5 | (Shiba et al., 2004) |

**Figure 1:**
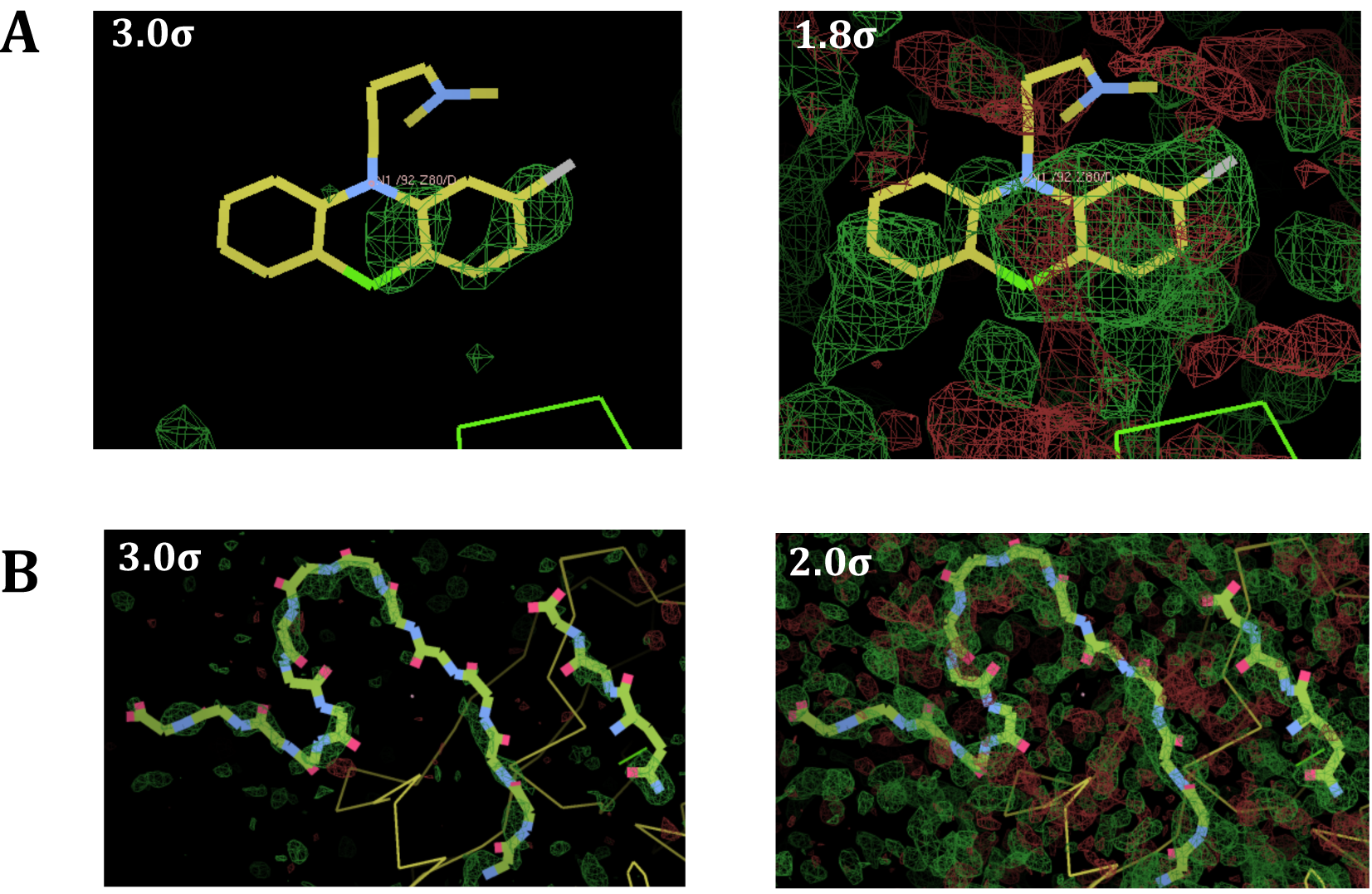
Delete refine *F*o-*F*c maps for representative structures. (A) Maps for one of the two ligands in the asymmetric unit in PDB: 3LK0 (Wilder et al., 2010), contoured at 3.0σ and 1.8σ and (B) maps for two peptides in the asymmetric unit in PDB: 2J9N (Viola et al., 2007) contoured at 3.0σ and 2.0σ. Positive and negative electron densities are shown in green and red colors, respectively. Maps were generated after deleting the ligands and refining the structures in PHENIX (Adams et al., 2010). Figures were generated using COOT (Emsley and Cowtan, 2004).

### B-Factors

Temperature factor or B-factor is an important parameter while monitoring the quality of model during the structure refinement. However, an appropriate theoretical model for refinement of B-factors that will enable universally acceptable refinement has not yet been possible, and the available strategies are constantly evolving (Merritt, 2012). The crystallographic refinement involving temperature factors (B individual vs. B group) ought to be based on resolution and the observation to parameter ratio rather than automated procedures. Refinement of B-factors of individual atoms is acceptable at high resolution, but the similar treatment of B-factors at lower resolution leads to artifacts (Merritt, 2012). The quality of data is required to be viewed in the context of specific problem under scrutiny and its functional implications. Indeed, depending on the resolution and the context, skilled crystallographers usually adopt appropriate strategies to avoid model bias.

Recently, it has been suggested that differences in the B-factors of the ligand and the protein imply that the ligand is not bound to the protein (Stanfield et al., 2016a). This contention is basically erroneous as there are a number of situations wherein the B-factors of the ligand and the protein can be dissimilar. One reason could be that the affinity of the ligand binding is relatively low due to which not all macromolecules in the crystal have ligand bound to them and this will be reflected in the B-factors. The B-factors would also be higher if the ligand does not have rigid geometry and the contacts with the protein are limited. The resolution of a number of structures of functionally important complexes, such as those of the helicase and primase (PDB: 2R6C) involved in DNA replication is low (~4 Å) and the atoms refine with very high B-factors (~150 Å) (Bailey et al., 2007). In comparison to the intrinsically flexible molecules such as linear peptides, rigid ligands are expected to have better defined electron density and much lower B-factors unless they bind in multiple conformations. It is anticipated that flexible ligands such as peptides may bind in more than one conformation for the backbone and the bound peptides may exhibit further variation in the side chain rotamers. To avoid constitutive activation/repression of corresponding biological processes, the majority of interactions between biological molecules have to be necessarily transient. The inherent dynamic nature of macromolecular interactions associated with physiological processes is often reflected even in static crystal structures. As a result, the application of excessively stringent parameters that ignore the biological context of the molecular interactions under scrutiny will render unnecessary rejection of the vast majority of protein-ligand complex structures.

Table 2 lists a few from the many examples to support the argument that B factors in the case of physiologically relevant bound ligands are higher than those of the protein. Most of the ligands cited here, which are either peptides or nucleic acids, exhibit higher degree of freedom in backbone conformation. Notably, due to variations in side chain conformations, peptides show even greater degree of freedom. This degree of freedom is considerably reduced when ligand binds protein with higher affinity through multiple sites of stabilizing interactions.

**Table 2:** Comparison of protein and ligand B-factors deposited in a representative structure in PDB.

| Sl. No | PDB code | B-factors as reported in deposited coordinates |  | Resolution | Reference |
| --- | --- | --- | --- | --- | --- |
| | | ($\text{\AA}^2$) | ($\text{\AA}^2$) | | |
|  |  | Protein | Ligand |  |  |
| 1 | 4CSA | 74.2 | 122.9 | 2.3 | (Leyrat et al., 2014) |
| 2 | 4MFL | 31.8 | 79.8 | 1.9 | (Lyu et al., 2014) |
| 3 | 4MFP | 38.2 | 83.9 | 2.2 | (Lyu et al., 2014) |
| 4 | 4N60 (Chain A) | 79.0 | 140.8 | 2.9 | (Xu et al., 2013) |
| 5 | 3KL5 | 58.3 | 94.9 | 2.6 | (St John et al., 2011) |
| 6 | 3LK0 | 43.5 | 113.5 | 2.0 | (Wilder et al., 2010) |
| 7 | 3LK1 | 49.8 | 91.3 | 1.8 | (Wilder et al., 2010) |
| 8 | 2J9N | 12.9 | 69.4 | 1.5 | (Viola et al., 2007) |
| 9 | 1UVI | 32.7 | 121.1 | 2.2 | (Salgado et al., 2004) |
| 10 | 1UVM | 43.0 | 112.6 | 2.0 | (Salgado et al., 2004) |
| 11 | 1PWW | 44.8 | 99.5 | 2.8 | (Turk et al., 2004) |
| 12 | 1HI6 | 45.1 | 79.5 | 2.6 | (Keitel et al., 1997) |

### Ramachandran Parameters

Ramachandran map has been a time-tested method of validating protein stereochemistry. Theoretically, except due to structural constraints, the protein model should abide by the limits set according to Ramachandran map. However, it is important to recognize that the structures deposited in the PDB are experimentally obtained data and could reflect typical constraints of such data. It has been established that the crystallographic refinement will be more sensitive to Ramachandran parameters at higher resolution as compared to that at lower resolution (Morris et al, 1992; Gunasekaran et al., 1996). During the early years, the PDB had a larger proportion of structures at low resolution as the data were often collected at home sources. Therefore, PROCHECK (Laskowski, 1993) which was based on the earlier PDB, was generous with respect to the allowed region of the Ramachandran map (Read et al., 2011). In the recent past, a large number of high resolution structures have been determined due to the availability of high intensity X-rays at synchrotron sources. This large database of high resolution structures is utilized by MolProbity (Chen et al., 2010), and this program is much more stringent in assessing stereochemical quality. Therefore, the structures that are at moderate resolution and were earlier assessed using PROCHECK, when subjected to MolProbity will naturally exhibit a large number of residues in the disallowed region. Conversely, it is also to be emphasized that good stereochemistry does not necessarily imply well-determined structure as stereochemistry could be guided by imposing tight constraints (Morris et al., 1992).

Analyses of high-resolution protein structures have shown that some residues can be present in the disallowed regions of the Ramachandran map (Gunasekaran et al., 1996; Morris et al., 1992). In some cases, these residues have been shown to be important for the function of the protein (Gunasekaran et al., 1996). It is possible that an individual phi, psi outlier is conformationally correct and arises due to biological/structural relevance in a given context. The presence of residues in the disallowed regions of the Ramachandran map is possible due to interactions formed between the backbone atoms of that residue with side chains of neighboring residues. The residue at the 51^st^ position in the light chain is present in the disallowed region in a number of antibody Fab structures (Nair et al., 2000).

It is observed that the resolution of the structure and the number of outliers are correlated (Morris et al., 1992). It is important to note that Ramachandran parameters couldn’t be directly correlated with presence/absence of the electron density, despite the emphasis to the contrary (Stanfield et al., 2016b). For low-resolution datasets, the corresponding structure could refine very well even if the stereochemistry is compromised. Therefore, a small fraction of residues is commonly observed in disallowed region of the Ramachandran map in the intermediate/low-resolution structures deposited in PDB.

## Protein Data Bank Depositions

The current policy of the Protein Data Bank to present the validation reports along with the data is very useful. The depositors are provided with these validation reports and have the opportunity of reviewing their structures before depositing. What is really appreciable is that the PDB does not sit on the judgment regarding the structures. It is left to the users to decide whether to use the data or otherwise, based on the quality parameters in the validation reports and the relevance to the biological problem under scrutiny. Many journals now ask for a submission for the report along with the manuscript to enable reviewers to use it along with electron density map figures in the manuscript to make an informed judgment about the quality and relevance of the structure to the major findings in the manuscript. Such a policy is perfectly in order and will be very useful, as the authors will have the opportunity to provide a rationale for the deviations, if any, with respect to the validation reports prior to the decision of acceptability or otherwise is made.

## Supporting Biochemical Evidence

Currently, structural data is almost always supported by appropriately designed biochemical experiments. Macromolecular folding leads to surface complementary which is not incorporated in refinement methods (Read et al., 2011). Therefore, biochemical methods assessing intra- and intermolecular interactions are useful for independent validation of structural observations. Importance of biological relevance in macromolecular crystallography is such that a number of structures are never deposited or documented that do not conform with relevant biochemical/genetic experiments.

While various validation mechanisms implemented by PDB ensure quality of the deposited structural data, the one-size-fits-all approach does not hold ground, particularly with challenging datasets wherein the ligand exhibits low occupancy or multiple conformations. Due to the diversity of ligand binding affinities, protein thermodynamics and crystallographic resolution, it is not prudent to apply benchmarks of uniform stringency on structural data to assess quality.

We would like to reinforce that while dealing with biological systems several variables are critical and therefore all cases should not be subjected to uniform criteria, which may be restrictive and arbitrary. Such criteria would result in limited applicability and immense loss of genuine structural data of high biological relevance. Considering the constraints that the macromolecular crystallographic experimentations impose, one has to carefully distinguish between signal and noise. It needs to be emphasized that structural biologists determine macromolecular structures to address specific questions in biology. The structure needs to be of a quality adequate enough to answer questions that are addressed in that particular study. Analyses using multiple parameters ought to be used, as is done by many experienced macromolecular crystallographers, some of whom are cited above. If one is to blindly follow restrictive criteria, a lot of biologically relevant structural data will go into oblivion. If this happens in case of drug-discovery studies, it would be particularly unfortunate. The erroneous strategy of seeking only high quality structural data for ligands may be especially lethal for fragment-based drug-discovery studies (Murray and Blundell, 2010). Application of automated and excessively stringent parameters will lead to unnecessary rejection of low-affinity binders that can be evolved through chemical modification to obtain binders with high affinity and specificity.

## Spurious Challenges

The deposited data corresponding to published works are sometimes abused to mount spurious challenges. In fact, work from our laboratory has been subjected to such attacks (Sethi et al., 2006; Khan and Salunke, 2012, Tapryal et al., 2013; Khan and Salunke, 2014). To understand how the humoral immune system generates enough receptors that have the cumulative capacity to recognize any and all non-self antigens, we had characterized the interaction of different antibodies with peptides unearthed using a phage display library (Manivel et al., 2000; Manivel et al., 2002). Our efforts showed that flexibility in terms of backbone and side chain conformation**s** in the peptide antigens and the antibody paratopes enables the germline antibodies to bind different antigens with adequate affinity. These studies involved structures of Fab fragments of antibodies with the low-affinity peptides, which were obtained through the refinement procedures outlined in the Introduction. These studies were questioned by a group of people who seem to be incapable of dealing with challenging datasets (Stanfield et al., 2016a&b). In spite of adequately responding to all the questions raised and the journal closing the subject by publishing their queries and our rebuttals (Salunke et al., 2016 a&b), the group did not stop misrepresenting our data in different ways.

It is not correct to post-facto apply arbitrary criteria for judging selectively the data which have been obtained using established methodologies and crystallographic experience and which have gone through due review process and are accepted based on the standard norms. It is acceptable if someone wants to analyze the entire PDB based on any new criteria. But cherry-picking specific structures with ulterior agenda and then cleverly misrepresenting the data, is by itself a fraud and needs to be stopped. The challengers appear to cleverly exploit the fact that it is relatively simple to carry out flawed refinement and generate electron density maps that lack clarity. The harmful intentions of these individuals can be gauged from the fact that their initial efforts to get the research papers retracted were all anonymous and when journal refused to accept their argument unilaterally, they were forced to go public with their point of view. It is unfortunate when people sitting on the judgment on other people’s data do not apply similar yardsticks to their own structures. This is evident when one re-analyses some of the published structures (Viola et al., 2007; Wilder et al., 2010).

It has been stated that the standard approach according to modern practice is to inspect the difference electron density omit map contoured at 3.0σ (Stanfield et al., 2016a). This is absolutely invalid. To justify 3.0σ cut-off for ligand fitting, the reasoning given subsequently (Weichenberger et al. 2017), is because it is the default contouring level in the display program Coot. To say the least, this is absolutely arbitrary. There ought to be more rational crystallographic/statistical logic for insisting on 3.0σ. It is a routine practice to vary the σ cutoff while interpreting electron density map; this is something every experienced crystallographer practices and reports. Use of delete-refine *F*o-*F*c maps at levels less than 3.0σ, often at 1.5σ contour level for visualizing the ligand model is routinely practiced (Table 1). In fact, the ligand model could not have been interpreted in the two structures (PDBs: 2J9N and 3LK0) without visualizing *F*o-*F*c electron density maps below 3σ cut-off as shown in Figure 1 (Viola et al., 2007; Wilder et al., 2010).

To state that the overall B factors of neighboring components in a crystal structure cannot differ drastically is shear hypocrisy (Stanfield et al., 2016a), considering that two of the co-authors have published structures where the B values of the ligands (PDBs: 2J9N and 3LK0, 3LK1 and 3KL5) are very high compared to neighboring amino acid residues of the proteins to which they bind (Viola et al., 2007; Wilder et al., 2010). Figure 1 shows that two of these structures (PDBs: 2J9N, 3LK0) when viewed at 3.0σ cutoff fail to show interpretable electron density in delete-refine *Fo-Fc* maps. It would be required to look for positive omit difference density at significantly lower σ cutoff (Figure 1) while the B-factors are relatively high (Table 2). It is strange that claims are made that the present validation methods are not adequate to judge correctness of data and yet certain structures are being selectively targeting by citing the validation reports based on the very same methods (Weichenberger et al. 2017). The reality is, validation tools are fine, but there are twilight zones where the judgment based on crystallographic experience is far more vital than the automated tools. The sound judgment and abilities of the experienced crystallographer take precedence over the application of excessively stringent automated criteria to prevent the loss of precious information of biological relevance.

It is important to note that we are dealing with not mature but germline antibodies or antibodies with polyspecificity which bind to the antigens with relatively lower affinity as the binding site is potentially flexible and therefore, as expected, antigen atoms exhibit higher B-factors (Khan and Salunke, 2012, 2014; Sethi et al., 2006; Tapryal et al., 2013). Germline antibodies possess unique thermodynamic properties that are physiologically relevant (Manivel et al., 2000). These thermodynamic properties are the result of flexibility in the germline antibody paratope, which results in an inherently lower binding affinity. Further, this flexibility is the basis for the cross-reactivity exhibited by germline antibodies, which is the primary step in antibody recognition of the antigenic universe. The situation in our structures is similar and is not unexpected given the biology of germline antibodies. The criticism of our data is unwarranted as the aforementioned researches have different and generous set of standards for their own structures but impose excessively high quality benchmarks that are incompatible with the resolution and biology on the existing norms practiced by other crystallographers in describing their structures.

The potential damage that spurious challenges can cause can be gauged from the situation involving structural studies with DNA polymerase iota (Nair et al., 2004). The investigators realized that the asymmetric unit contains two molecules of the enzyme bound to each end of the substrate DNA and this entire complex occupies two conformations-related by a 180° inversion in the crystal. Gradual model building and crystallographic refinement using this description of the asymmetric unit led to electron density maps that showed the presence of a Hoogsteen base pair in the active site (Nair et al., 2004). This finding was challenged by an individual who chose to describe the contents of the asymmetric unit differently and subjected the data to “black-box” refinement and this exercise led to maps that showed no clear density in the active site (Wang, 2005). In the above example, subsequent biochemical and structural studies showed unambiguously that the initial finding was correct and above reproach (Nair et al., 2005; Johnson et al. 2005; Johnson et al., 2006; Nair et al., 2006). The entire episode highlights the strength of approach described in the Introduction and illustrates how an important finding could have been consigned to obscurity due to erroneous description of contents of the asymmetric unit and excessively hasty refinement.

Ramachandran map is indeed an excellent yardstick for quality of protein structures. Yet, a detailed analysis of the PDB shows that there exist plenty of structures in the PDB that violate the Ramachandran parameters beyond generally accepted limits. There are more than 200 structures in PDB with around 10% or greater Ramachandran outliers. Table 3 shows a list of randomly selected structures among them. For example, Duffy-binding-like domain from *Plasmodium knowlesi* (PDB: 2C6J) and the nucleosome assembly protein S from *Plasmodium falciparum* (PDB: 3KYP) show Ramachandran outliers at the level of 10% and 15%, respectively (Gill et al., 2010; Singh et al., 2006). Also, structure of a nitrogenease from *Azotobacter vinelandii* (PDB: 1NIP) exhibits Ramachandran outliers as high as 12% (Georgiadis et al., 1992). (Wlodawer et al. 1989) have deposited in PDB a structure with Ramachandran outliers above 10% (Table 3). These represent much higher level of Ramachandran outlier than in any structures from our laboratory (Khan and Salunke, 2012, Khan and Salunke, 2014; Sethi et al., 2006; Tapryal et al., 2013).

**Table 3.** Statistics of Ramachandran outliers in some randomly selected entries from PDB.

| **Sl No** | **PDB code** | **Resolution (Å)** | **Ramachandran Outlier (%)** | **Reference** |
| --- | --- | --- | --- | --- |
| 1 | 3KYP | 2.8 | 15 | (Gill et al., 2010) |
| 2 | 2C6J | 3.0 | 10 | (Singh et al., 2006) |
| 3 | 1UK0 | 3.0 | 14 | (Kinoshita et al., 2004) |
| 4 | 1M6E | 3.0 | 13 | (Zubieta et al., 2003) |
| 5 | 1MUO | 2.9 | 13 | (Cheetham et al., 2002) |
| 6 | 1M5N | 2.9 | 12 | (Cingolani et al., 2002) |
| 7 | 1G4A | 3.0 | 13 | (Wang et al., 2001) |
| 8 | 1JDE | 2.8 | 10 | (Ye et al., 2001) |
| 9 | 2DL2 | 3.0 | 12 | (Snyder et al., 1999) |
| 10 | 4PRG | 2.9 | 11 | (Oberfield et al., 1999) |
| 11 | 1NIP | 2.9 | 12 | (Georgiadis et al., 1992) |
| 12 | 3HVP | 2.8 | 11 | (Wlodawer et al., 1989) |

A view point article (Rupp et al., 2016) has been published essentially to criticize policies of the PDB and the journal editors for not giving in to the one sided coercion. A recent publication also continues to harp on the same issues under the guise of describing a new version of software (Weichenberger et al., 2017). In science, if there is a dispute about interpretation of data or about the conclusions made in a paper, it is by no means scientific misconduct or fraud. We believe that the journal editors follow normal practice by responding to the comments received and seeking explanations from the authors on the papers published in their journals. In some cases, the editors decide to publish both the letters and the rebuttals and in the other cases the editors reject the spurious challenges based on the responses of the authors. In either case, due process of scientific debate is followed. History is full of examples where there have been strong views on the results of others, sometimes proved valid and at other times not. In this specific matter, it is sad to see some colleagues conflate disagreement however strong with misconduct and then to follow this up by stirring a witch-hunt. The journal editors, reviewers and the PDB understand that the final judgment regarding quality of macromolecular structures is largely dependent on the context of the biological problem under scrutiny.

While it requires enormous skills and experience to obtain biochemically meaningful interpretations from macromolecular crystallographic data, it is extremely easy to carry out erroneous refinement and generate fuzzy maps to create unnecessary and invalid controversies regarding deposited structures. It is also clear that structural information, for example, regarding flexible ligands that bind to macromolecules with micromolar affinity cannot be subjected to the same criteria as that of nanomolar affinity protein-ligand complexes. Such sweepingly uniform default approaches will lead to unnecessary rejection at the initial stages itself of potentially useful data that may otherwise contain important discoveries causing enormous damage to the field. Left to the vigilante groups, many important researches with implications to public health and biotechnology would be stillborn. In addition, the actions of such people result in journal editors becoming unnecessarily suspicious of structural papers and this will lead to several disastrous consequences, the worst being that it will drive many young scientists away from utilizing the method of macromolecular crystallography.

## Acknowledgements

I would like to thank my colleagues and friends, many of whom are accomplished macromolecular crystallographers, for help and discussions.

